# Mucin-mimetic action of capsaicin improves high fat diet-induced gut barrier dysfunction in mice colon

**DOI:** 10.1101/2021.08.04.455153

**Authors:** Vijay Kumar, Vibhu Kumar, Neha Mahajan, Jasleen Kaur, Kirti Devi, Ravinder Naik Dharavath, Ravindra Pal Singh, Kanthi Kiran Kondepudi, Mahendra Bishnoi

**Author notes:** **<u>Corresponding authors</u> Mahendra Bishnoi, PhD**, Scientist E, TR (i)P for Health Laboratory, Division of Food and Nutritional Biotechnology, National Agri-Food Biotechnology Institute (NABI), Knowledge City-Sector 81, SAS Nagar, Punjab 140306, India., **KanthiKiran Kondepudi, PhD**, Scientist E, |Healthy Gut Research Group, Division of Food and Nutritional Biotechnology, National Agri-Food Biotechnology Institute (NABI), Knowledge City-Sector 81, SAS Nagar, Punjab 140306, India.

## Abstract

The gut barrier – including tight junction proteins and mucus layers, is the first line of defense against physical, chemical, or pathogenic incursions. This barrier is compromised in various health disorders. Capsaicin, a dietary agonist of Transient receptor potential vanilloid 1 (TRPV1) channel, is reported to alleviate the complications of obesity. While its mode of action is well established to enhance energy expenditure, metabolism and prevent dysbiosis, the more local effects on the host gut – particularly the gut barrier and mucus system remain elusive.

We employed a diet-induced obesity model to investigate the effect of capsaicin on the gut barrier and mucus production and to understand the involvement of mucus, bacteria, and TRPV1 in these phenomena. Mucin feeding reflected most of the effects produced by capsaicin, indicating that mucus modulation by capsaicin plays a crucial role in its anti-obesity effects. Capsaicin, bacteria and the host mucus system seem to act in a cyclic cascade involving TRPV1, which can be activated by capsaicin and various bacteria. These findings provide new insight into the role of TRPV1 in maintaining a healthy gut environment.

**HIGHLIGHTS:**

- Exogenous mucin feeding produced anti-obesity effects similar to capsaicin in mice.
- Mucin and capsaicin improved TJP expression, intestinal permeability and gut microbial diversity.
- Capsaicin modulated bacterial diversity *in vitro*, independently of the host.
- Probiotic bacteria and butyrate activated TRPV1 in transfected HEK cells.
- Anti-obesity action of capsaicin is not exclusive to TRPV1 agonism.
- Capsaicin’s action as colonic mucus secretagogue – plays a crucial part in its anti-obesity benefits.

## 1. INTRODUCTION

Mucus acts as the first line of defense in the gastrointestinal tract against physical insults and pathogenic invasions. Mucus in the gut is arranged in two layers – inner mucus, tightly bound to epithelial surface and generally impermeable to bacteria; and outer mucus, which is loosely bound and exposed towards gut lumen [1]. The building blocks of mucus are highly glycosylated proteins called mucins. Mucins are produced and secreted by specialized cells called goblet cells, present in intestinal crypts as cell clusters. More than 20 types of mucins have been characterized, generally categorized into two types – trans-membrane (contribute to cellular signaling and immune responses) and gel-forming [2]. Mucus forms the physiological barrier in the gut, along with tight junction proteins (TJPs) – zona occludens, occludin and claudins present in epithelial cell layers, preventing bacteria and harmful chemicals in the gut from entering the gut circulation [3]. Besides, the outer mucus layer harbours various bacterial populations, which interact with the host through different mechanisms, creating a healthy gut environment [4]. The mucosal lining in the gut is adversely affected in various diseases, including inflammatory bowel disease (IBD), colitis and metabolic disorders like obesity [5,6]. The primary reason for the deterioration of the mucosal barrier due to high-fat diet is the deficiency of complex carbohydrates in the diet and microbiota-mediated degradation of mucus layer in the gut. Gut dysbiosis is also linked to the impaired gut barrier function and compromised permeability, leading to endotoxemia and subsequent dysfunctions in metabolism and immune responses [5,7].

Transient receptor potential vanilloid 1 (TRPV1) is a non-selective cation channel, which transports Ca^2+^ ions upon activation by a variety of stimuli, such as heat (temperatures >43°C), acidic pH and several dietary molecules – capsaicin (chilli), allyl isothiocyanate (mustard), gingerol (ginger), piperine (pepper), etc. TRPV1 is primarily present on sensory neurons and acts as a nociceptor, inducing pain and inflammatory responses upon activation. It has been extensively studied in relation to inflammation, immune response, carcinogenesis and nociception in the past [8,9]. In recent years, the expression of TRPV1 in the gastrointestinal tract, its dietary modulation and consequent beneficial effects have been a focus of research. The role of TRPV1 in preventing gastrointestinal complications like colitis and IBD and metabolic disorders such as obesity and type II diabetes is explored using various agonists/blockers [10,11]. Capsaicin, an active ingredient of chilli, is the most studied agonist of TRPV1. In numerous reports, capsaicin administration prevented high fat diet-induced weight gain, improvde glucose homeostasis and energy expenditure, and altered gut microbiota profile with increased abundances of healthy bacteria [12-17]. Most of the research on capsaicin and obesity is focused on its role in energy expenditure, thermogenesis, glucose metabolism, insulin sensitivity, lipid accumulation/breakdown, inflammation, feeding behavior, etc [12-17]. While multiple researchers studied the effects of capsaicin on gut microbiota, its role as a mucin secretagogue in exerting the anti-obesity effects, remains elusive.

Some reports relate capsaicin to mucus secretion in the respiratory tract and the role of TRPV1 in increased expression of mucin genes *MUC2, MUC5AC* [18,19]. Mucous secretion is reported to be regulated by inflammatory cytokines, neurotransmitters and hormones, which interestingly are also known to be affected by TRPV1 modulation [20,21]. TRPV1-positive neurons profusely innervate the gut, including intestines, and mediate responses to stimuli via the local release of neurotransmitters and/or by communicating to the central nervous system. In a novel finding from our previous study, resiniferatoxin-induced TRPV1 ablation in rats led to a drastic decrease in mucus production and dysbiosis in the colon [22]. In the present study, we used a diet-induced obesity model in mice, studied alterations in gut barrier function, and explored what role CAP, TRPV1, gut microbiota, and particularly mucus itself might play in alleviating high fat diet-induced impairments.

## 2. METHODS

### 2.1. Animals

Male C57BL6/J mice (aged 4-6 weeks) were procured from IMTech Centre for Animal Resources and Experimentation (iCARE), Chandigarh, India, and housed in Animal Experimentation facility at National Agri-Food Biotechnology Institute (NABI), Mohali, India. Animals were kept in a pathogen-free environment at 25±2°C and maintained on a 12-hour light-dark cycle. After 1 week of acclimatization, they were randomized and divided into 4 groups (n=8 each) – Control (normal pellet diet-fed), HFD (high fat diet-fed; ∼60% energy from fat), H+MUC (HFD + 1g/kg/day porcine stomach mucin p.o.) and H+CAP (HFD+ 2mg/kg/day capsaicin p.o.). All animals were given free access to water and food. The interventions were given for 12 weeks, after which the animals were sacrificed and relevant samples were harvested for further studies. Animals were weighed weekly and upon sacrifice, weights of white adipose tissue (WAT) and liver were recorded. Institutional Animal Ethics Committee (IAEC) of NABI approved experimental protocol (Approval number NABI/2039/CPCSEA/IAEC/2020/05). The experiment was conducted according to the Committee for the Purpose of Control and Supervision on Experiments on Animals (CPCSEA) guidelines on the use and care of experimental animals. A brief plan of the experiment is shown in Figure 1.

**Fig. 1.**
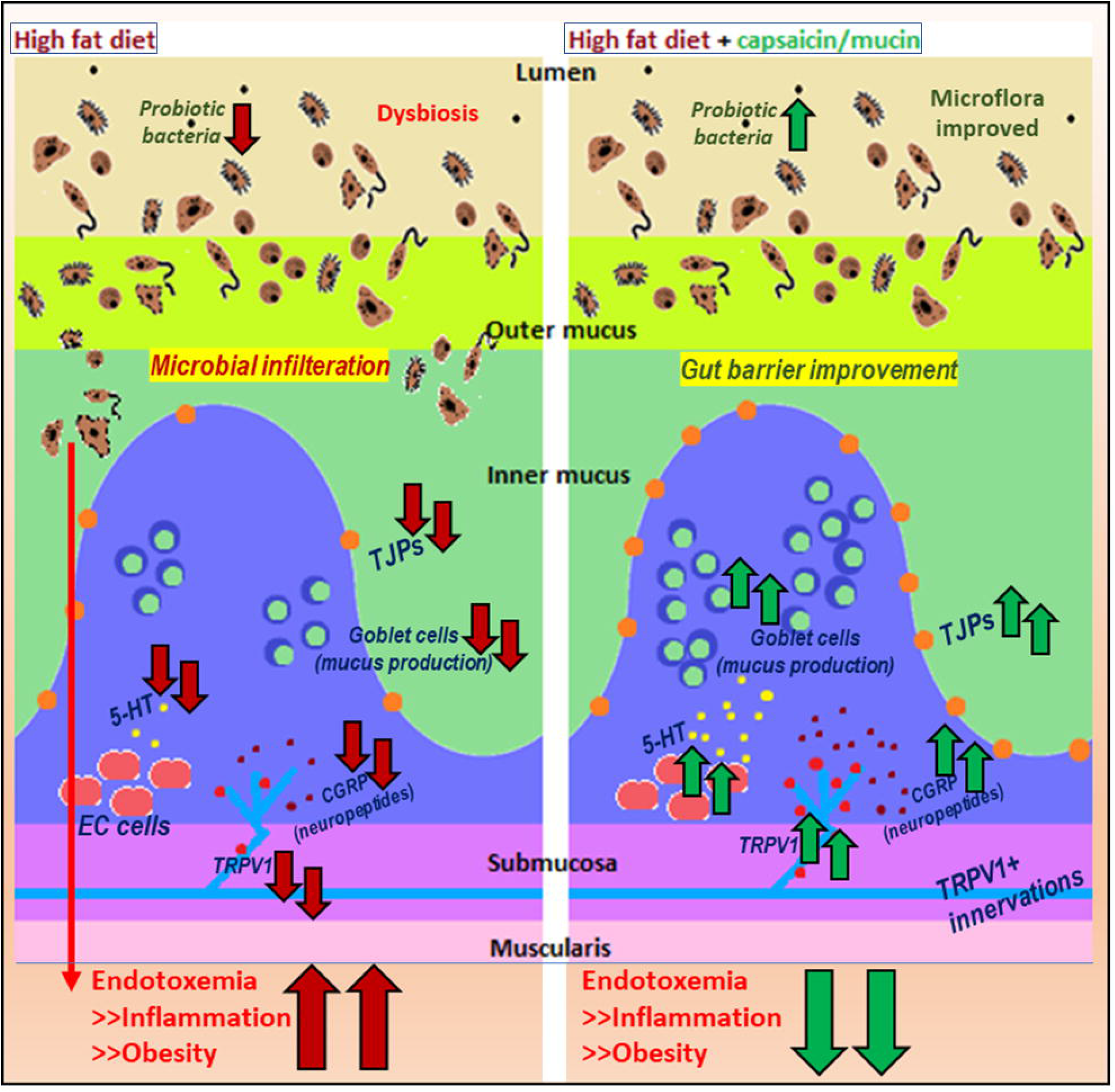
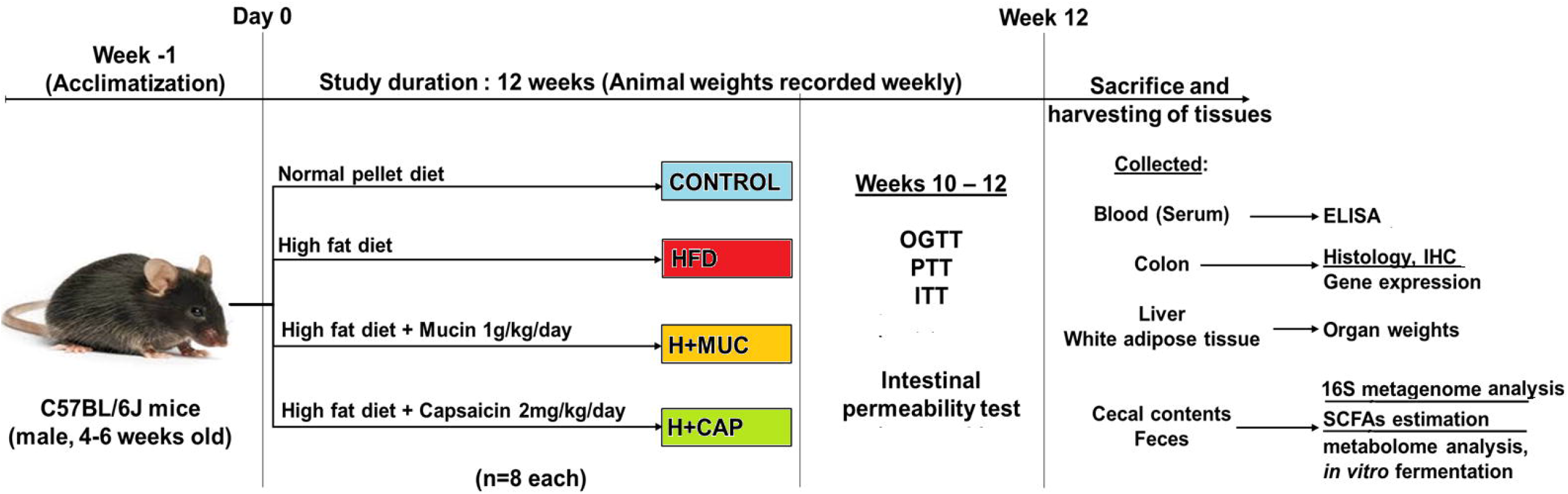
Animal experiment plan.

### 2.2. Diet and interventions

A normal pellet diet was procured from Hyalasco Biotechnology Pvt Ltd (India). A high-fat diet (HFD; 60% energy from fat) was prepared in-house, with slight modifications in a previously described composition [23]. The composition of HFD was: Normal pellet diet (36.5% w/w), Milk casein (25.0% w/w; Loba Chemie, India); Animal lard (35.0% w/w; local market, Chandigarh, India), Vitamin and mineral mix (6.0% w/w), DL-methionine (0.3% w/w), Sodium chloride (0.1% w/w; Himedia Laboratories Pvt Ltd, India). Capsaicin (≥95%, from *Capsicum sp*.) and Porcine Stomach Mucin (Type III, bound sialic acid 0.5-1.5 %) were purchased from Sigma-Aldrich (USA).

### 2.3. Blood glucose measurement

Oral glucose tolerance test (OGTT) and insulin tolerance test (ITT) were done during the 11^th^ and 12^th^ weeks of study to assess glucose homeostasis. Animals were fasted for 12h and 6h, for OGTT and ITT, respectively. A 2g/kg body weight p.o. Glucose; and 1U/kg i.p. insulin were used for these tests. Blood glucose levels were measured at 0 (before treatment), 15, 30, 60, 90 and 120min after treatment, using Glucocard (Arkray, Japan) glucometer.

### 2.4. Intestinal permeability test (*in vivo* imaging and serum FITC-dextran estimation)

Intestinal permeability tests were performed during the 11^th^ and 12^th^ weeks of the study. 600 mg/kg body weight Fluorescein isothiocyanate–dextran (average mol wt. 3,000-5,000, Sigma-Aldrich, Missouri, USA) in 0.9% saline was given to mice orally. Animals were subjected to hair removal and fluorescence imaging (IVIS Lumina Series III, Perkin Elmer, USA) of the abdominal region at 0 (before treatment), 30, 60, 90 and 120 min after treatment, under anesthesia (Isoflurane: induction at 4%, maintenance at 2.5-3%). Blood was collected before and 90min after treatment, and serum was separated by centrifugation at 8000rpm for 10min. Fluorescence in serum was measured using excitation at 485nm and emission at 530nm (SpectraMax m5e fluorescent microplate reader, Molecular Devices, USA).

### 2.5. Histology and immunohistochemistry

After sacrifice, colon samples were stored in either 10% formalin (in PBS) or Carnoy’s fixative (60% ethanol, 30% chloroform, 10% glacial acetic acid) for further processing. For paraffin embedding, tissues were prepared by serial dehydration in ethanol (25%, 50%, 70%, 90%, 100%, 2 h each) followed by xylene treatment (2h, twice). After embedding (in molten paraffin at 60°C, 2 h, twice) and microtomy blocks formation, 5µm thick sections were made.

For staining, sections were de-paraffinized using xylene (2-5 min, twice) and subjected to serial rehydration in ethanol (100% to 25%, 2min each, followed into water). Alcian blue (AB)-Hematoxylin-Eosin(H&E) (Hi-media Laboratories, India) stains were used for 15 mins, 30 s, 15 s, respectively, with washing (thrice in water for 5min each, after every staining step). For sulfomucins staining, a High iron diamine (HID) stain was used for 24h, and Eosin was used as counterstain. Slides were then again serially dehydrated and treated in xylene and mounted using DPX mountant (Hi-media Lab, India). Tissue morphology and mucin production were observed in FFPE samples under 20X objective magnification (Leica CTR6, Leica Biosystems, Germany), and mucus layer thickness was estimated in Carnoy’s fixative-fixed tissues. Staining intensity and mucus layer thickness were quantified using ImageJ software (NIH, USA).

For studying the expression of proteins – Zona occludens-1, Claudin-1, Occludin and TRPV1, immunohistochemistry was performed in FFPE sections. After de-paraffinization and serial rehydration, sections were subjected to antigen retrieval (in citrate buffer pH 6.0, at 90°C for 30min) and blocking (5% goat serum in PBS-T, Tween-20 0.1%). Primary antibodies - Zona occludens 1 (ZO1; 200X dilution in 2.5% goat serum, Cat. No. 13663S, Cell Signaling Technology, USA), Occludin (200X dilution, Cat. No. 91131S, Cell Signaling Technology, USA), Claudin 1 (200X dilution, Cat. No. 91131S, Cell Signaling Technology, USA) and TRPV1 (100X dilution, Cat. No. NBP1-97417, Novus Biologicals, LLC, USA) were used overnight in a humidified chamber at 4°C. After washing in PBS-T (3 times, 5 min each), secondary antibody Alexa fluor 488 (1000X dilution, Cat. No. 4412S, Cell Signaling Technology, USA) was used for 2h at room temperature. 1µg/ml PureBlu DAPI (Biorad Laboratories, USA) was used as counterstain. Imaging was done using confocal microscopy (Carl Zeiss LSM-880, Carl Zeiss, Germany). The analysis for antibody fluorescence intensity was done using ImageJ software (NIH, the USA).

### 2.6. Bacterial abundance

16S rRNA V3-V4 region based metagenome analysis was done in cecal content. For this purpose, the bacterial genomic DNA was isolated from the cecal content of mice using the NucleoSpin DNA Stool kit (Macherey Nagel, Düren, Germany), following the manufacturer’s instructions. It was used in 16S metagenome analysis on the HiSeq Illumina sequencing platform (outsourced to NxGen Bio Diagnostics Pvt. Ltd., New Delhi, India). Raw reads quality assessment (threshold value Q20, 99.9% reads qualified), paired reads assembly, OTUs processing and microbiome analysis were done using online available resources (https://rdp.cme.msu.edu, http://cgenome.net/calypso).

### 2.7. Short-chain fatty acids (SCFAs) estimation

SCFAs in the cecal content were measured using HPLC (Agilent 1260 Infinity series chromatographic system; Agilent Technologies, Singapore) by following a previously described method [24]. Briefly, 100-120 mg of sample was added to 500 µl of acidified water (pH 2) and homogenized by thorough vortexing followed by a 10 min incubation at room temperature before centrifugation (6000rpm/20 min/4°C). The supernatants were filtered using Millex-GN 0.2 µm nylon syringe filters (Millipore, Massachusetts, USA). A 20 µl sample was injected into the Hi-Plex H column (300×7.7 mm; 8 µm particle size, Agilent Tech.). 0.1% formic acid in Milli-Q water (Merck Millipore, 0.22 µm filtered, resistivity 18.1–18.3 MΩ cm) was used as mobile phase. The column was equilibrated and eluted with an isocratic flow rate of 0.6 ml/min at 50 °C for 60 min. Volatile Acids Mix (Cayman Chemicals Co., USA) was used as standard at concentration range 100-6400µM. Final sample concentrations were calculated as µM/mg.

### 2.8. Serum ELISAs

Insulin, leptin, LPS and IL-6 levels were estimated in serum collected after sacrifice. Respective ELISA kits (insulin, leptin, IL-6 – MilliPlex Map Mouse Adipokine kit, Merck, Germany; LPS – E2214Mo, Bioassay Technology Laboratory, UK) were used as per the manufacturer’s instructions. Colorimetric readings were taken using SpectraMax m5e fluorescent microplate reader (Molecular Devices, USA) or MAGPIX multiplex system (Merck, Germany).

### 2.9. Serotonin estimation

Serotonin levels were measured using HPLC in serum and colon. Samples were homogenized in ice-cold acetonitrile: water (50:50) solution containing 0.1% formic acid and centrifuged at 15000rpm for 15min at 4°C. The supernatant was filtered through 0.2 μm nylon filters (MDI Advanced Microdevices Pvt Ltd, India) before injecting into a reverse-phase octadecyl silane (C18) HPLC column (YMC Triart 250mm×4.6mm, ID S-5 120 A, 5µm, YMC Co. Ltd, Japan). HPLC system with an electrochemical detector (Waters 2465, Waters Corporation, USA) was used. The mobile phase consisted of acetonitrile: water (25:75) solution containing 0.1% formic acid at a flow rate of 0.9mL/min and pH-3.0 with a run time of 10 min. The retention time for serotonin peaks was 5.8min. Data obtained was analyzed by Empower Pro software (Waters Corporation, USA).

### 2.10. Fecal *in vitro* fermentation

Fresh feces were collected from normal pellet diet-fed C57BL/6J mice (n=12) and pooled into 3 samples. The slurry was made by dissolving 100 mg feces in 1 ml PBS, which was further diluted 10-fold in Basal medium (total volume=10ml each) supplemented with glucose and without/with capsaicin. Microbiological media of the following composition was used: peptone water 2g/l, yeast extract 2g/l, NaCl 0.1g/l, K_2_HPO_4_ 0.04g/l, KH_2_PO_4_ 0.04g/l, MgSO_4_.7H_2_O 0.01g/l, CaCl_2_.6H_2_O 0.01g/l, NaHCO_3_ 2g/l, Tween-80 2ml, hemin 0.02g/l, vitamin K1 10μl, cysteine-HCl 0.5g/l, pH 7.0). Our *in vivo* dose of 2 mg/kg in 1ml/kg volume of the vehicle corresponds to ∼655 µM concentration. Literature suggested that 85-95% of orally given capsaicin is absorbed within 1-2h of administration [25]. This leaves approximately 60 µM capsaicin available for chronic interaction with gut bacteria. We used two concentrations – 50 µM and 100 µM for our experiment. Control, CAP50 and CAP100 (n=4 each) groups were assigned to treatments and fermentation was carried out for 48h at 37°C in anaerobic conditions. After 48h, culture samples were pelleted down at 8000rpm for 20min. The pellets were subjected to bacterial DNA isolation (see section 2.6) and qPCR was performed to evaluate bacterial abundance. Method details for qPCR (Supplementary methods: b) and the list of primers used (Table 1) is given in supplementary data.

### 2.11. HEK-293 cells transfection and Ca_2+_-influx measurement

HEK-293 cell line was procured from National Centre for Cell Science, Pune, India. Cells were maintained in DMEM (Lonza, Switzerland) containing 10% FBS (Lonza, Switzerland) and 1X PenStrep (Thermo Fisher Scientific, USA) at 37°C, 5% CO_2_. All experiments were performed between passage numbers 25-35. Plasmid pVQ CMV NanoV1-2a-EGFP ferritin was purchased from Addgene, USA (Addgene plasmid #79649, deposited by Jeffrey Friedman) [26]. Cells were seeded in 96 well plates, and at 70-80% confluency, transfection was carried out using Lipofectamine 3000 Reagent (Thermo Fisher Scientific, USA), as per supplier’s instructions (0.2µg DNA/well, 0.4µl transfection reagent/well). After 36-48h, transfection was verified by GFP visualization and TRPV1 activity (through Ca^2+^-influx measurement) was confirmed using capsaicin (50µM). Fura-2AM (Thermo Fisher Scientific, USA) fluorescent dye was used for intracellular Ca^2+^ measurements. HEK cells were incubated in a medium containing 10µM Fura-2AM for 45min at 37°C. Then, media was removed and respective treatments were given (n=6-8 each). Immediately, the cells were subjected to fluorimetry (SpectraMax m5e fluorescent microplate reader, Molecular Devices, USA) at excitation 340 nm and 380 nm, emission at 510 nm, through 2 min at 20s intervals). F340/F380 ratio was interpreted as intracellular Ca^2+^ levels.

### 2.12. Activation of TRPV1 by selected putative probiotic strains

*Lacticaseibacillus rhamnosus* (Lab3), *Lactiplantibacillus plantarum* (Lab39) and *Bifidobacterium breve* (Bif11) strains were isolated and characterized at Healthy Gut Research Group, National Agri-Food Biotechnology Institute (Bif11: [27], Lab3/39: *unpublished data*). *L. rhamnosus* and *L. plantarum* strains were grown in MRS under aerobic conditions and *B. breve* was grown in MRS supplemented with L-cysteine hydrochloride (0.05%, w/v) under anaerobic conditions (Anoxomat Mark II, Mart Microbiology BV, Netherlands) at 37°C for 48 h. Butyrate (2.5mM; Sigma Aldrich, USA), live Lab3, Lab39 and Bif11 cultures (48h culture in 10ml culture media, diluted to OD 1.0 (∼10^10^ CFU/ml), pelleted, washed and further diluted in 5ml PBS with Ca^2+^) were evaluated for TRPV1-agonistic action.

### 2.13. Statistics

Data were analysed using GraphPad Prism 8 software (GraphPad, USA). All data are presented as mean±S.E.M. (bar graphs, line graphs), individual values (dot plots) or only average values (heatmaps). One-way ANOVA with Bonferroni’s test was employed to assess the differences between the groups for single parameters. Two-way ANOVA with Tukey’s multiple comparisons was used in the case of repeated measurements/multiple parameters. P<0.05 was considered significant in all data (however, a numerical p-value is also given at certain places to include some major but statistically insignificant changes).

## 3. RESULTS

### 3.1. Capsaicin and mucin prevented HFD-induced metabolic complications in mice

HFD feeding caused a significant increase in animal weights, which was prevented in H+MUC and H+CAP groups (Fig.2A). Organ weights (liver and WAT) were also significantly higher in the HFD group, while they were decreased in both interventions (Fig.2B). OGTT test revealed significantly higher glucose intolerance in the HFD group at almost all time points. On the other hand, mucin and capsaicin administration prevented this change. AUC analysis reflected a significant improvement in the H+CAP group compared to HFD. The difference was observable but not statistically significant in H+MUC (Fig.2C). ITT also showed the same pattern. After 30 min of insulin administration, the blood glucose clearance was significantly less in HFD, while both interventions improved (Fig.2D-E). Serum insulin levels were significantly higher in the HFD than Control group, whereas both the interventions significantly prevented this alteration (Fig.2F). Serum leptin levels were significantly higher than Control in the HFD group. Mucin and capsaicin prevented it; however, the difference was not statistically significant (Fig.2G).

**Fig.2.**
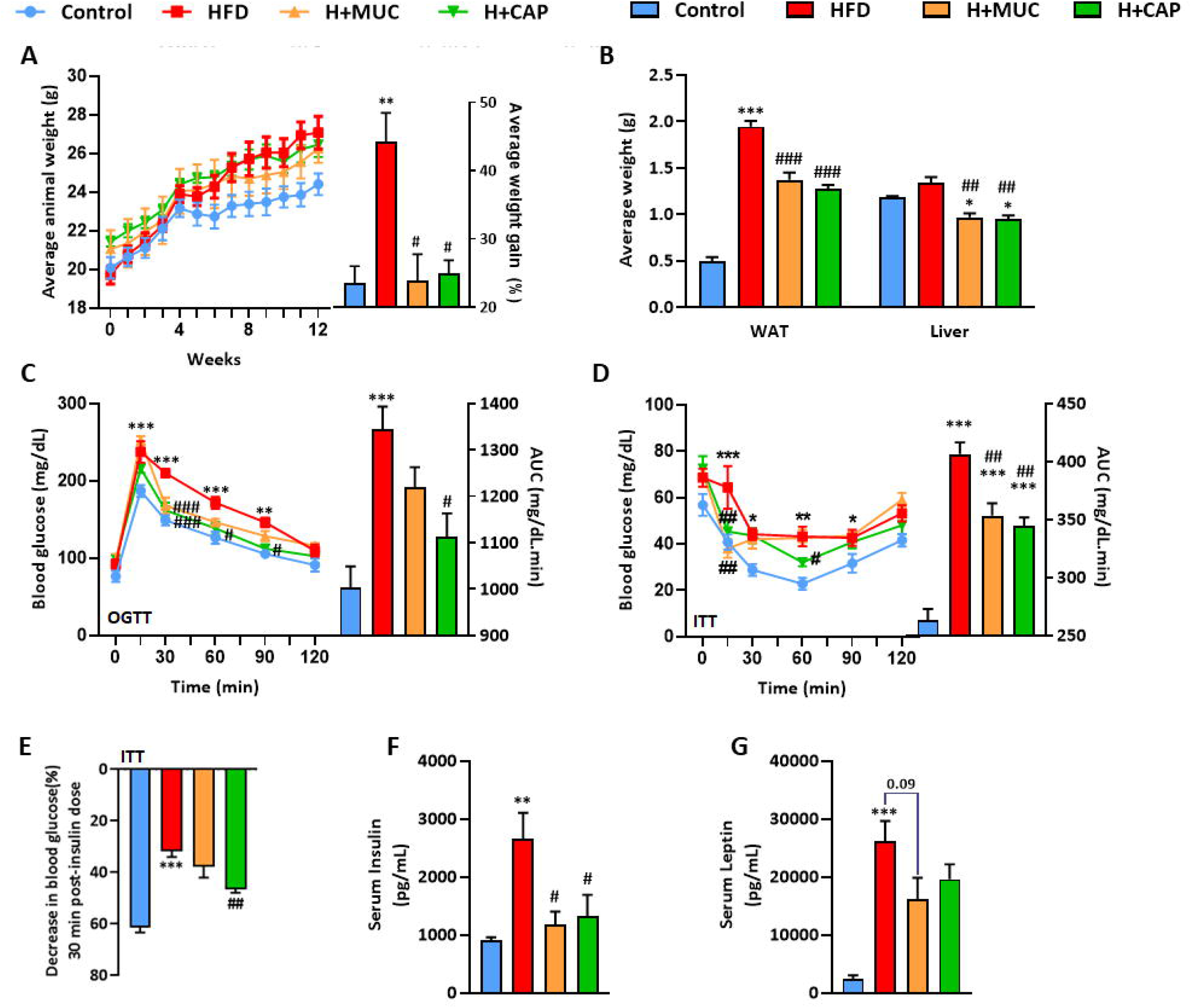
Effect of high fat diet and interventions on metabolic parameters. A) Weekly animal weights and total % weight gain, B) Organ weights, C) OGTT – glucose measurements and AUC, D) ITT – glucose measurements and AUC, E) % decrease in glucose levels upon insulin administration, F) Serum insulin and G) Serum leptin levels. Mice were divided into 4 groups (n=8 each) – Control (normal pellet diet-fed), HFD (high fat diet-fed), H+MUC (HFD + porcine stomach mucin (1g/kg/day p.o.), H+CAP (HFD + capsaicin 2mg/kg/day p.o.). Food and water were given *ad libitum*. Treatments were given for 12 weeks. Animal weights were recorded weekly (n=8 each). Oral glucose (2g/kg p.o.) tolerance test (OGTT) and Insulin (1IU/lg i.p.) tolerance test (ITT), were performed during week 10 and 11 (n=6 each). Upon sacrifice, organs weights were recorded (n=8 each) and Serum ELISAs were performed (n=5 each). All data is represented as mean ± SEM. Intergroup variations were assessed using One-way ANOVA with Bonferroni’s multiple comparisons test (% changes, AUCs, organ weights, insulin, leptin) or Two-way ANOVA (repeated measures) with Tukey’s multiple comparisons test (blood glucose levels). *p<0.05, **p<0.01, *** p<0.001 versus Control; #p<0.05, ##p<0.01, ###p<0.001 versus HFD.

### 3.2. Capsaicin and mucin prevented HFD-induced increase in permeability, endotoxemia and inflammation markers, and improved gut barrier function

*In vivo* fluorescent imaging showed a major decrease in mean fluorescent intensity in the target abdominal area with time in the HFD group, indicating higher diffusion rates of FITC-dextran into circulation through the gut barrier. In H+MUC and H+CAP groups, the magnitude remained comparable to Control throughout the experiment (Fig.3A-B). Serum fluorescence measurement in 90min samples also reflected the same phenomenon, with a significantly higher fluorescence level in the HFD group. In contrast, both interventions significantly prevented it, with levels similar to Control (Fig.3C). Serum LPS (endotoxemia) and IL-6 (inflammation) levels were significantly higher in the HFD group, while they were significantly reduced in both interventions (Fig.3D-E).

**Fig.3.**
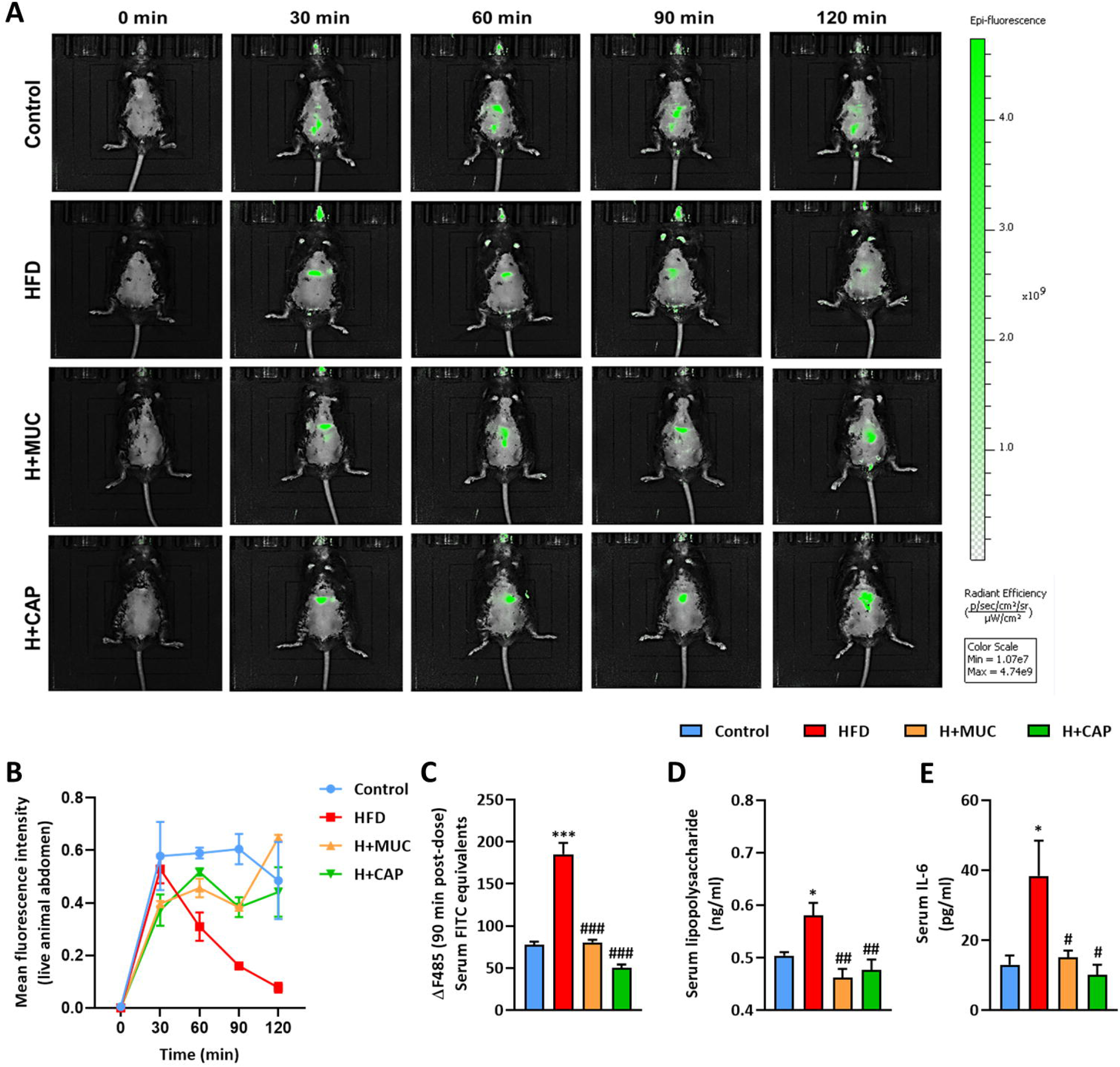
Effect of high fat diet and interventions on intestinal permeability, endotoxemia and inflammation. A) FITC-dextran *In vivo* fluorescent imaging – representative figure, B) FITC fluorescence intensity quantification in abdominal region at different time points, and C) Serum FITC fluorescence measurement at 90min, D) Serum LPS and E) Serum IL-6 levels. Mice were divided into 4 groups – Control (normal pellet diet-fed), HFD (high fat diet-fed), H+MUC (HFD + porcine stomach mucin (1g/kg/day p.o.), H+CAP (HFD + capsaicin 2mg/kg/day p.o.). Food and water were given *ad libitum*. Treatments were given for 12 weeks. During week 11 and 12, intestinal permeability was tested. Single dose (n=3 each) of FITC-dextran (600mg/kg p.o., mol. wt. 3-5kDa) was given, animals were anaesthetized for live imaging at 0 (before treatment), 30, 60, 90 and 120 min after treatment. *In vivo* fluorescence intensity (n=3 each) was measured using ImageJ software, serum FITC levels were measured at 480nm (ex.) - 520nm (em.). After sacrifice, serum LPS (n=6 each) and IL-6 (n=5 each) were measured using commercial kits. All data is represented as mean ± SEM. Intergroup variations were assessed using One-way ANOVA with Bonferroni’s multiple comparisons test. *p<0.05, **p<0.01, *** p<0.001 versus Control; #p<0.05, ##p<0.01, ###p<0.001 versus HFD.

Immunohistochemical analysis revealed that the expression of TJPs – ZO-1, Claudin-1, and Occludin were significantly diminished in HFD group sections compared to Control, indicating compromised gut barrier function. Both interventions showed improved expression of these proteins. ZO1 and Claudin1 had more improvement in the H+CAP group compared to H+MUC. Occludin expression was found equally improved by both interventions. TRPV1 expression was reduced in the HFD group and capsaicin administration increased it; however, the effect of HFD on TRPV1 expression was found unaffected by mucin treatment (Fig.4A-B).

**Fig.4.**
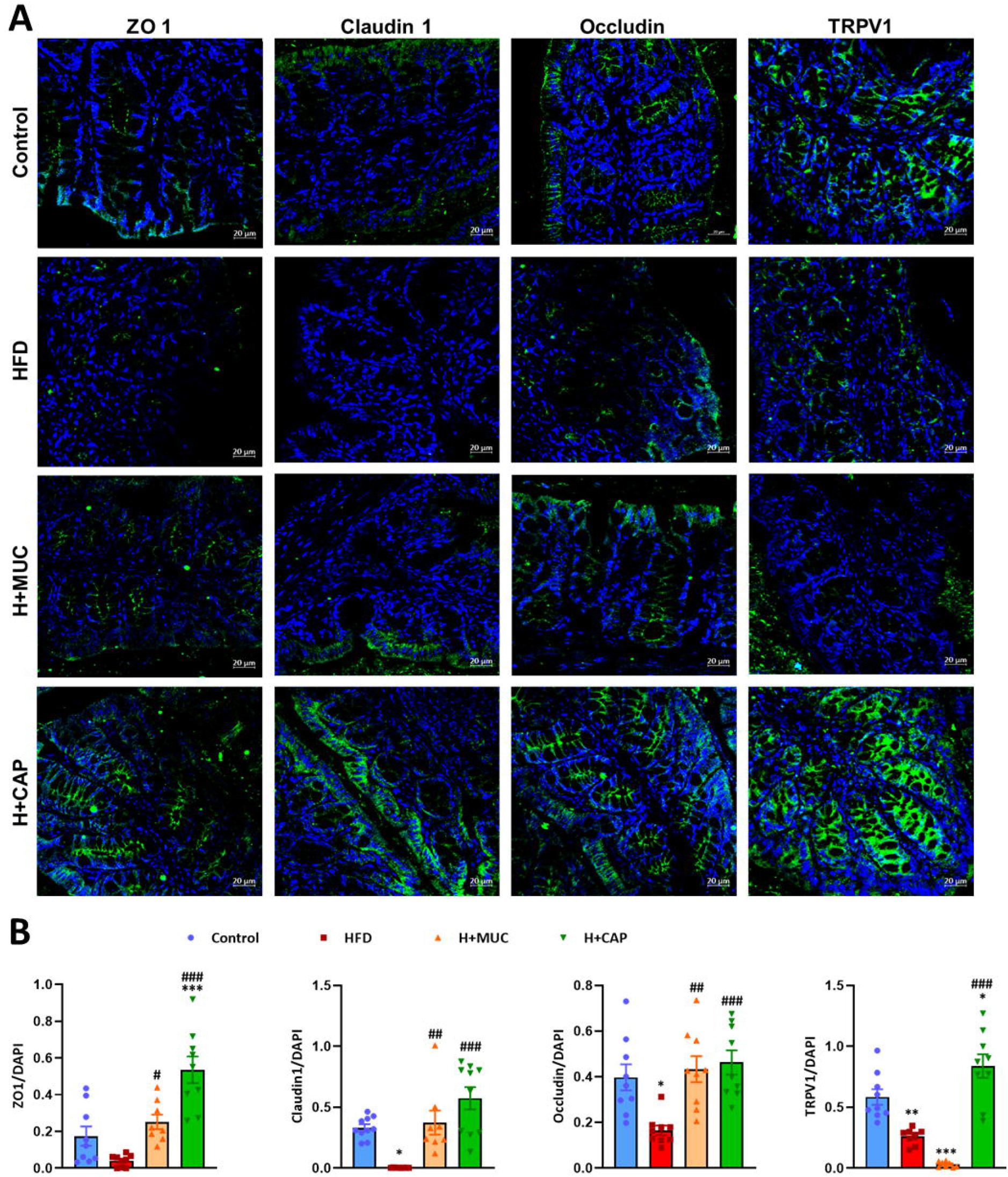
Effect of high fat diet and interventions on tight junction proteins expression and TRPV1. A) Immunohistochemistry (fluorescence) – representative images of Zona occludens 1 (ZO1), Claudin 1, Occludin and TRPV1, B) Fluorescence intensity analysis (antibody signal normalized to DAPI) Mice were divided into 4 groups – Control (normal pellet diet-fed), HFD (high fat diet-fed), H+MUC (HFD + porcine stomach mucin (1g/kg/day p.o.), H+CAP (HFD + capsaicin 2mg/kg/day p.o.). Food and water were given *ad libitum*. Treatments were given for 12 weeks. After sacrifice, colon tissues (n=3 each) were formalin-fixed and subjected to IHC (F) [antigen retrieval: citrate buffer (pH6, 90°C, 30 min), blocking: 5% goat serum in PBS-Tween20 (0.5%), incubations: primary antibody - overnight, secondary (FITC) antibody - 2h, DAPI - 2min]. Imaging was done by confocal microscopy and intensity analysis was done using ImageJ software (n=9 each; 3 measurements per sample image). All data is represented as mean ± SEM. Intergroup variations were assessed using One-way ANOVA with Bonferroni’s multiple comparisons test. *p<0.05, **p<0.01, *** p<0.001 versus Control; #p<0.05, ##p<0.01, ###p<0.001 versus HFD.

### 3.3. Capsaicin and mucin prevented HFD-induced alterations in colonic mucus production

AB and HID stains were used to study colonic morphology and mucus production (Fig.5A, C). ImageJ analysis of intensity (normalized to whole section area) of AB stain revealed that total mucin staining in the HFD group was significantly lower than Control. H+MUC and H+CAP groups showed levels of AB uptake similar to Control, indicating a preventive effect of both interventions on high fat diet-induced alterations in mucus production ((Fig.5B). HID stain, specific for sulfomucins, showed an opposite pattern, where the HFD group showed higher HID uptake than Control. Both interventions reversed this effect (Fig.5D).

**Fig.5.**
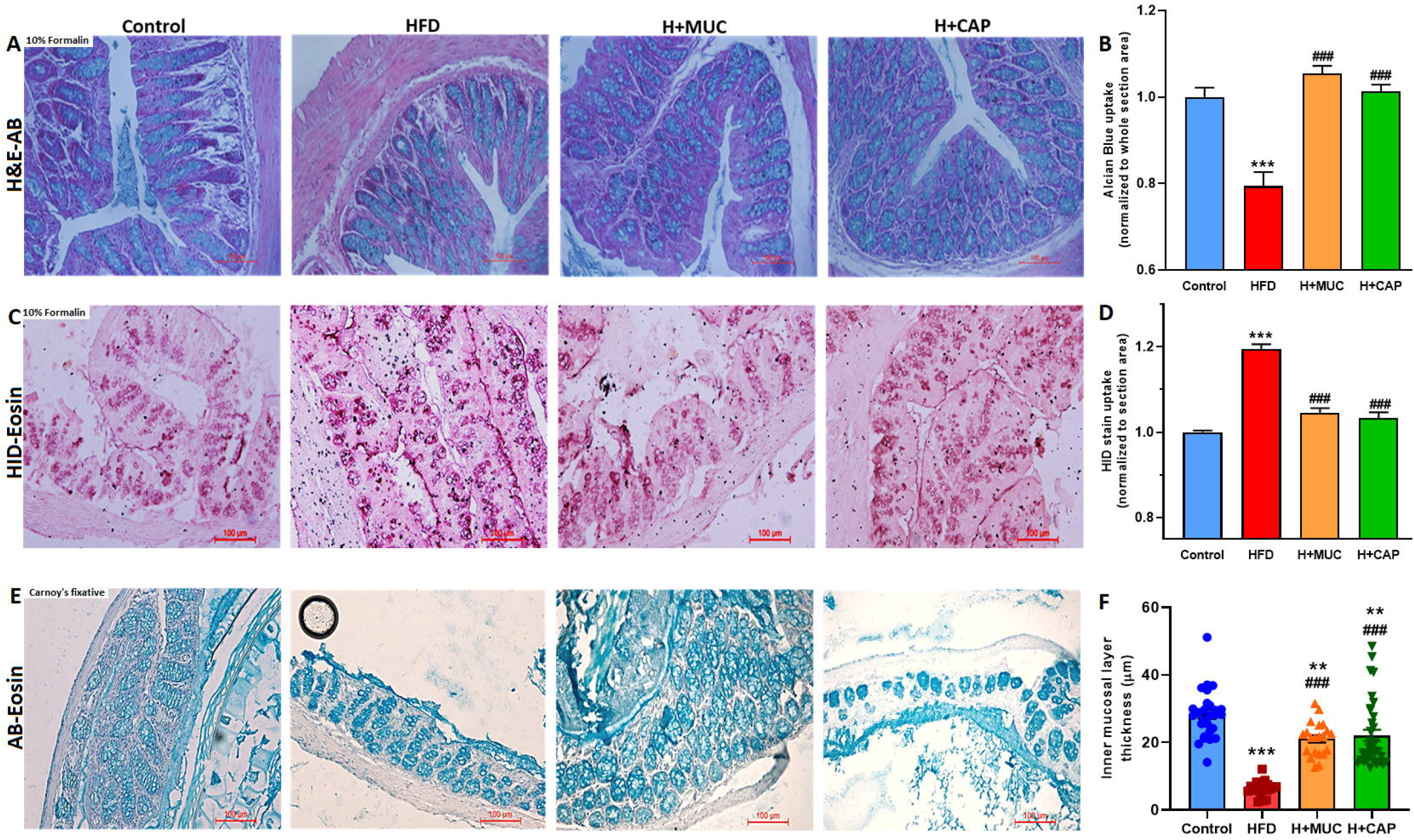
Effect of high fat diet and interventions on colonic mucus production. Histology – representative images of A) Hematoxylin-Eosin-Alcian blue (H&E-AB) staining and B) Alcian blue intensity, C) High iron diamine-Eosin (HID-Eosin) staining and D) HID staining intensity, E) Alcian blue-Eosin (AB-Eosin) staining (in Carnoy-fixed sections) for inner mucus and F) Average inner mucus layer thickness. Mice were divided into 4 groups – Control (normal pellet diet-fed), HFD (high fat diet-fed), H+MUC (HFD + porcine stomach mucin (1g/kg/day p.o.), H+CAP (HFD + capsaicin 2mg/kg/day p.o.). Food and water were given *ad libitum*. Treatments were given for 12 weeks. After sacrifice, colon tissues were fixed (formalin or Carnoy’s fixative), subjected to serial dehydration (ethanol for 2h each: 25, 50, 70, 90%, twice 100%), xylene treatment (1h twice) and paraffin embedding. 5µm sections were cut. For staining, paraffin was removed using xylene (5min twice), sections were serially rehydrated (ethanol: 100% twice, 90, 70, 50, 25% and PBS, 2min each) and treated with respective stains (H&E: 2min each, AB: 15min, HID: 24h) followed by washing (5min, thrice after each stain) in water. After serial dehydration and xylene treatment, sections were mounted using DPX medium. Microscopy was performed for imaging. Respective staining intensities (normalized to counter-stained whole section area) and mucus layer thickness were measured using ImageJ software. All data is represented as mean ± SEM. Intergroup variations were assessed using One-way ANOVA with Bonferroni’s multiple comparisons test. **p<0.01, *** p<0.001 versus Control; ###p<0.001 versus HFD.

The thickness of the inner mucus layer was measured in AB stained sections obtained from Carnoy’s fixed colon samples (Fig.5E). The average thickness was found significantly lower in the HFD group compared to Control. While average thickness in both H+MUC and H+CAP was also substantially less than Control, both showed remarkable improvement compared to HFD (Fig.5F).

### 3.4. Mucin caused changes in gut microbiota profile similar to capsaicin

16S rRNA-based metagenomic analysis was performed in the cecal content of mice to examine the changes in gut microbiota profile. Heatmap (Fig.6A) presents an overview of relative changes in abundance of bacterial genera in different groups. The Bray-Curtis ANOSIM test revealed that the overall composition of total microflora at the genus level varied most between Control and HFD, with H+MUC and H+CAP having Bray-Curtis coefficientsbetween the former two groups (p=0.001, R=0.674). Principal component analysis of the dissimilarity matrix shows clustering of different groups (Fig.6B, C). The components in consideration (PC1:39%, PC2:25%) indicate that the observed pattern was not representative of whole populations, again attributed to a limited number of replicates and high variations. The clustering patterns effectively differentiated the Control group from others. Negative R-values corresponding to HFD and H+CAP indicate that intragroup variations were increased in these groups.

**Fig.6.**
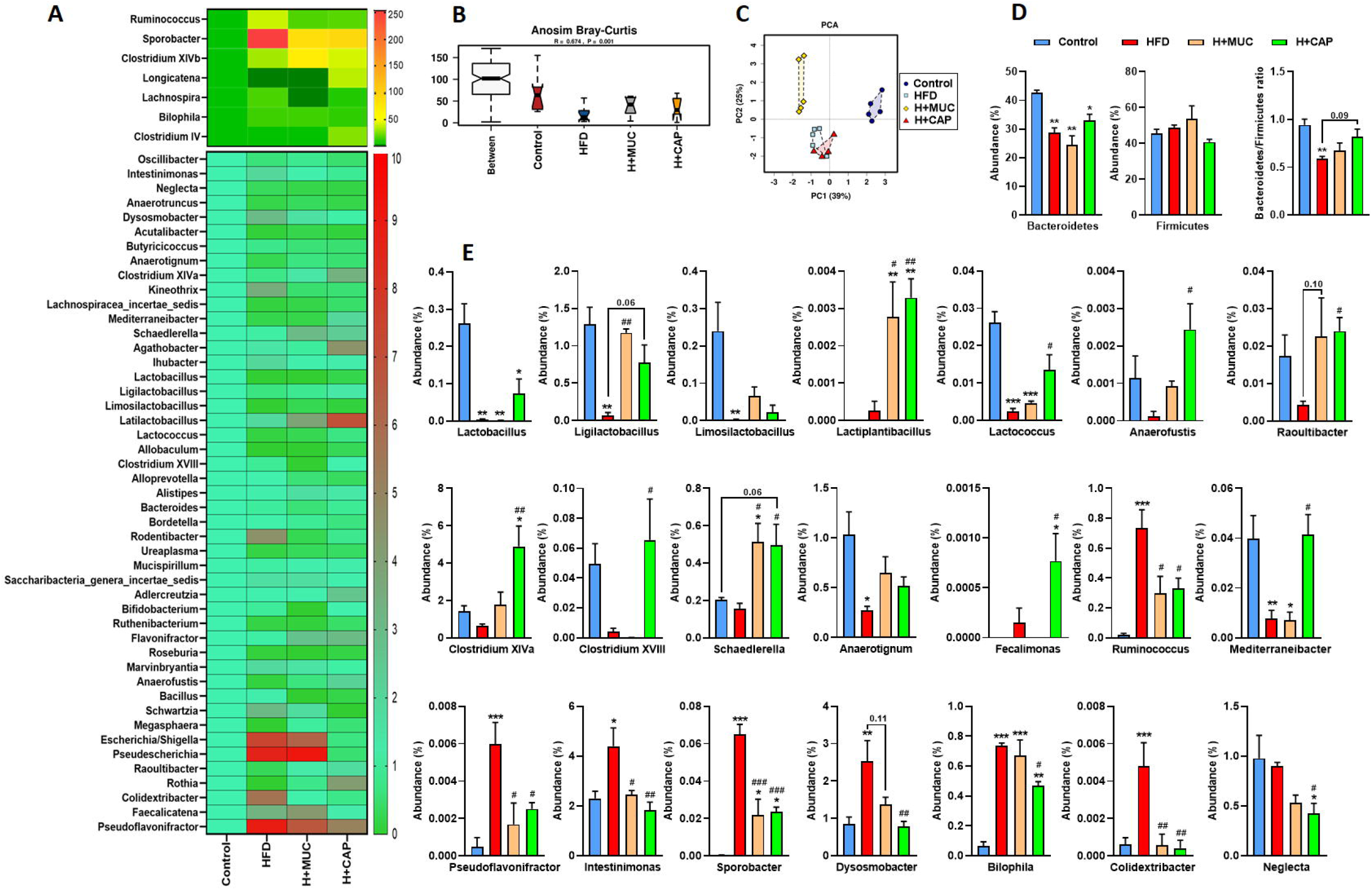
Effect of high fat diet and interventions on gut microbial population. A) Heatmap overview of bacterial abundance (genera), B) Bray-Curtis ANOSIM (dissimilarity index) and C) PCA plot, D) Abundance (%) of dominant phyla – Bacteroidetes and Firmicutes, and their ratio, E) Abundance (%) of various relevant bacterial genera in different groups. Mice were divided into 4 groups – Control (normal pellet diet-fed), HFD (high fat diet-fed), H+MUC (HFD + porcine stomach mucin (1g/kg/day p.o.), H+CAP (HFD + capsaicin 2mg/kg/day p.o.). Food and water were given *ad libitum*. Treatments were given for 12 weeks. After sacrifice, cecal contents were collected and bacterial genomic DNA was isolated (using commercial kit). DNA was subjected to paired-end Illumina sequencing for 16S rRNA variable region. Metagenome data was analyzed using online available resources (Calypso, RDPipeline). All data is represented as mean ± SEM. Intergroup variations were assessed using One-way ANOVA with Bonferroni’s multiple comparisons test. **p<0.01, *** p<0.001 versus Control; ###p<0.001 versus HFD.

The abundance of phylum Bacteroidetes and Bacteroidetes-Firmicutes ratio were found significantly lower in HFD compared to Control. Both intervention groups showed a ratio higher than HFD; however, the differences were not significant (HFD vs H+CAP; p=0.09) (Fig.6D). Among genera (Fig.6E), various lactic acid bacteria – *Lactobacillus, Ligilactobacillus, Limosilactobacillus*, and *Lactococcus* were decreased in the HFD group. Both interventions prominently improved the abundance of these genera. Another genus Lactiplantibacillus was increased significantly by both interventions. *Schaedlerella, Anaerofustis, Anaerotignum, Fecalimonas, Mediterraneibacter* and *Raoultibacter* were abundant bacteria from various *Clostridium* clusters negatively affected in HFD, which was reversed in both interventions. Genera like *Ruminococcus, Pseudoflavonifractor, Intestinimonas, Sporobacter, Dysosmobacter Bilophila* and *Colidextribacter* were significantly increased in HFD. Both interventions could mostly reverse the changes in their abundance. *Neglecta* genus was decreased in H+MUC and H+CAP.

### 3.5. Capsaicin modulated gut microbiota independently of host gut environment

Fresh feces from Control mice were subjected to fermentation in the presence of capsaicin to determine host-independent effects of capsaicin on gut microbes. qPCR results showed that 50µM dose of capsaicin increased Bacteroidetes-Firmicutes ratio and abundance of genera like *Bifidobacterium, Lactobacillus, Akkermansia, Eubacteria*, and *Fecalibacterium*. However, a higher dose (100 µM) did not produce these effects. The abundance of various gram-negative genera – *Vibrio, Prevotella, Bacteroides, Fusobacterium* and *Cronobacter*, remained unaltered at both doses (Fig.7A).

**Fig.7.**
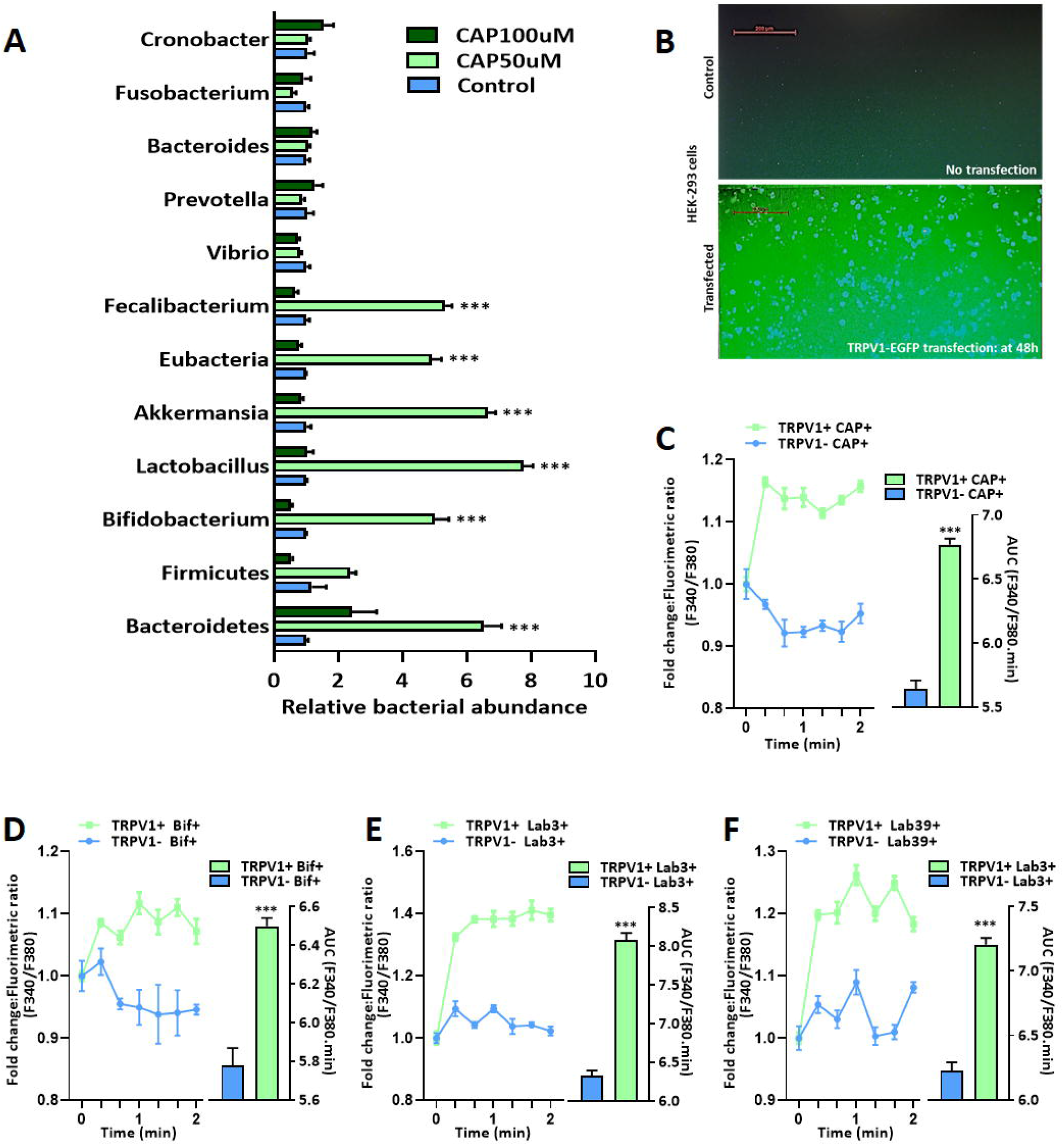
Interactions between capsaicin-gut bacteria-TRPV1 (*in vitro*). A) Relative abundance of fecal bacteria (after 48h fermentation without or with capsaicin 50µM/100µM), B) HEK cells transfection with TRPV1-EGFP – representative image, C-F) TRPV1^+^-HEK cells treatment - Fold changes in fluorometric ratio (representing Ca^2+^ influx) and AUCs (representing intracellular Ca^2+^ levels) upon various treatments [capsaicin 50µM (CAP), live *Bifidobacterium* (Bif11) and *Lactobacillus* (Lab3&39)]. For fecal fermentation, fresh feces were obtained from 15 normal C57BL6/J mice and pooled into a slurry of 0.1g/ml in PBS. 12 cultures were prepared by slurry dilution (10X) in culture media. Control and treatment groups - CAP50µM and CAP100µM (n=4 each) were incubated at 37°C under anaerobic conditions. After 48h, bacteria were pelleted and DNA was isolated. qPCR was performed using primers for various bacterial genera (16S rRNA). All data is represented as mean ± SEM. Intergroup variations were assessed using One-way ANOVA with Bonferroni’s multiple comparisons test. ***p<0.001 versus Control. HEK cells were transfected using lipofectamine, with gene for TRPV1-EGFP fusion protein. Transfection was visually confirmed by GFP imaging. TRPV1 activity was confirmed using Fura2AM dye. After 45min incubation with Fura2AM (10µM), cells were washed and treated with respective reagent/bacteria. Fluorescence at 340nm and 380nm was recorded (em. 510nm) every 20s for 2min. F340/F380 ratio was calculated and represented as fold change between TRPV1^-^(normal) and TRPV1^+^(transfected) cells. All data is represented as mean ± SEM. Intergroup variations were assessed t-test with Welch’s correction. *** p<0.001 versus Control.

### 3.6. Putative probiotic strains and butyric acid increased Ca^2+^-influx in TRPV1^+^ HEK cells

Transfection in HEK cells with a plasmid carrying the EGFP-TRPV1 fused gene was established by GFP visualization (Fig.7B). Intracellular Ca^2+^ measurement using Fura 2-AM dye showed that treatment with capsaicin caused Ca^2+^ influx into transfected cells, which confirmed the functioning of TRPV1 protein expressed by the gene insert (Fig.7C). Cells treated with live cultures of Lab3, Lab39 and Bif11 also showed significantly increased Ca^2+^ influx into TRPV1-transfected cells, as compared to normal (Fig.7D-F). A major bacterial metabolite, butyric acid, could elicit TRPV1-dependent Ca^2+^ influx (Fig.8D).

**Fig.8.**
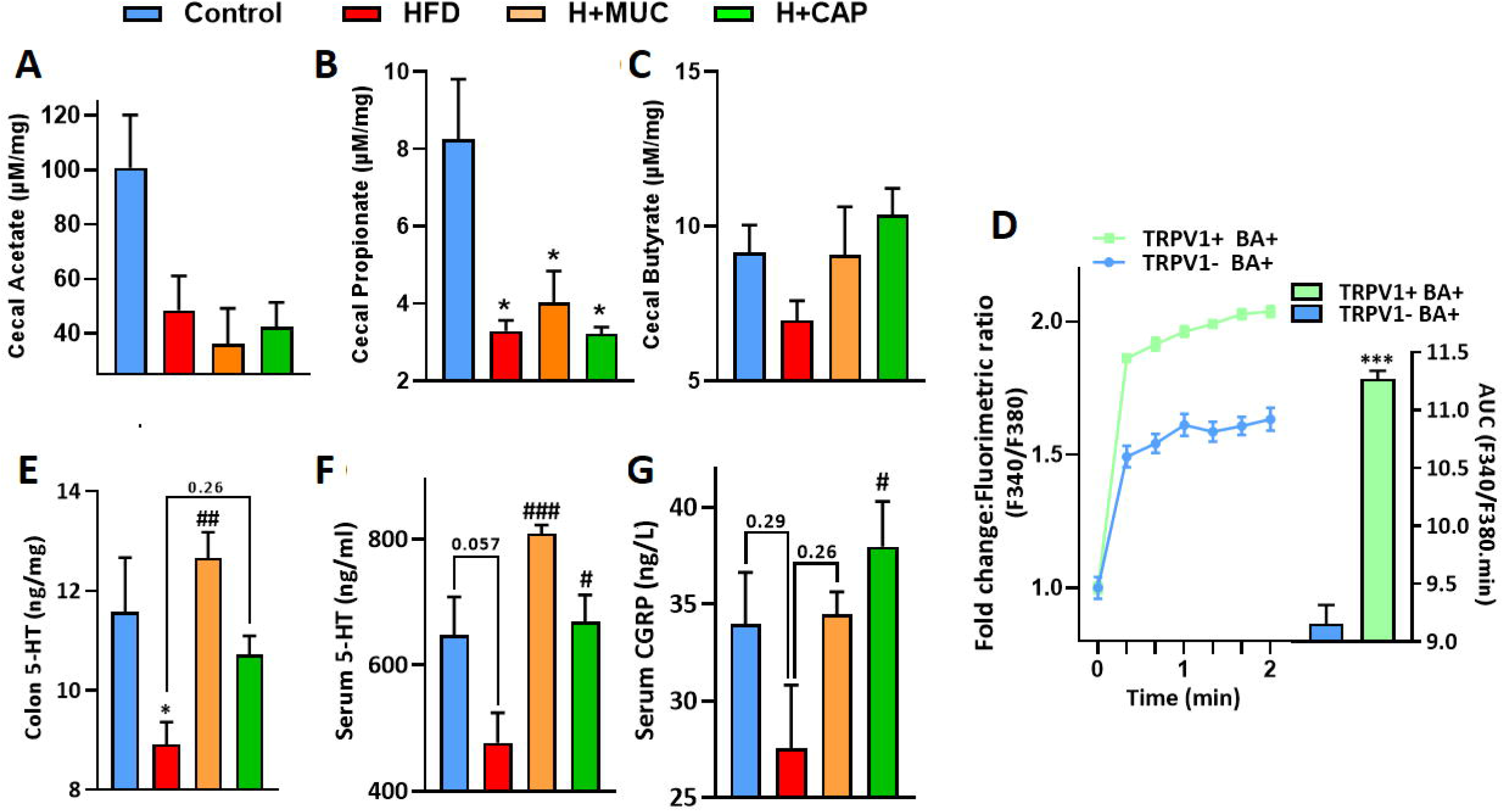
A-C) Major SCFA levels in cecal content, D) TRPV1^+^-HEK cells treatment - Fold change in fluorometric ratio (representing Ca^2+^ influx) and AUC (representing intracellular Ca^2+^ level) upon butyrate (BA 2.5mM) treatment, E-F) Colon and serum serotonin (5-HT) levels and G) serum calcitonin gene-related peptide (CGRP) levels. Mice were divided into 4 groups – Control (normal pellet diet-fed), HFD (high fat diet-fed), H+MUC (HFD + porcine stomach mucin (1g/kg/day p.o.), H+CAP (HFD + capsaicin 2mg/kg/day p.o.). Food and water were given *ad libitum*. Treatments were given for 12 weeks. After sacrifice, serum and colon samples were collected, homogenized, filtered and used in HPLC for Serotonin (5-HT) estimation [column: C18 reverse phase, mobile phase: 0.1% formic acid in water:acetonitrile: :50:50, detector: ECD]. Serum CGRP was estimated using ELISA kit. All data is represented as mean ± SEM. Intergroup variations were assessed using One-way ANOVA with Bonferroni’s multiple comparisons test. *p<0.05 versus Control; #p<0.05, ##p<0.01, ###p<0.001 versus HFD. HEK cells were transfected using lipofectamine, with gene for TRPV1-EGFP fusion protein. Transfection was visually confirmed by GFP imaging. TRPV1 activity was confirmed using Fura2AM dye. After 45min incubation with Fura2AM (10µM), cells were washed and treated with respective reagent/bacteria. Fluorescence at 340nm and 380nm was recorded (em. 510nm) every 20s for 2min. F340/F380 ratio was calculated and represented as fold change between TRPV1^-^(normal) and TRPV1^+^(transfected) cells. All data is represented as mean ± SEM. Intergroup variations were assessed t-test with Welch’s correction. *** p<0.001 versus Control.

### 3.7. Capsaicin and mucin reversed HFD-induced decrease in cecal butyrate, serotonin and calcitonin gene-related peptide (CGRP)

HPLC analysis of cecal content showed that the HFD decreased the levels of major SCFAs (acetate, propionate and butyrate) compared to Control (Fig.8A-C). Mucin and capsaicin interventions could increase butyrate levels. However, the changes were not found statistically significant. HPLC analysis of serum and colon showed that 5-HT levels were lower in the HFD group in both samples compared to Control. Both capsaicin and mucin improved 5-HT levels (Fig.8D, E). Serum CGRP also showed a similar pattern. HFD-induced decrease in CGRP was reversed by interventions, though only capsaicin showed significant improvement (Fig.8F).

## DISCUSSION

A high-fat diet is implicated in damaging the gut mucosa, as deficiency of dietary fibre leads to mucus foraging by bacteria [5,7]. We administered exogenous mucin to HFD-fed mice and compared its effects with the anti-obesity effects of capsaicin. Various characteristic parameters of obesity – weight gain, increased gross liver and white adipose tissue mass and impaired glucose metabolism were improved by mucin intervention, similar to capsaicin. Mucin also improved serum leptin, insulin and insulin resistance. We found a decrease in colonic mucus content and expression of tight junction proteins by HFD, while both capsaicin and mucin could restore them. These results suggested that increased availability of mucin/mucus in the intestines could be due to capsaicin supplementation, which might contribute to capsaicin’s anti-obesity effects.

FITC-dextran is used as a fluorescent reporter of gut permeability, with higher serum concentrations reflecting enhanced permeability and a compromised barrier function [28]. As expected, HFD group animals showed a steady decrease in fluorescence during live abdominal imaging. The high intensity was observed in serum samples, which pointed towards the HFD-induced “leaky gut.” Both capsaicin and mucin were able to prevent it significantly. For further examination, we used colon tissue in IHC. Major TJPs – ZO-1/2, Claudins and Occludin play an essential role in enforcing the gut barrier. The expression of these proteins was reported to be impaired in conditions such as obesity [29]. We also studied the expression of ZO-1, Claudin-1 and Occludin in colon sections. All of them were significantly reduced by HFD, whereas capsaicin expectedly increased their expression. Mucin also improved the expression of TJPs. Mucin itself is not assimilated by the digestive system of mice but acts as a nutrient for commensal bacteria in the gut and may improve the microbial diversity [30,31]. A number of bacteria, including probiotics like Lactobacillus and Bifidobacterium, enhance TJPs expression through their metabolites [32,33]. While capsaicin itself has been reported to cause the opening of TJs and enhance permeability [31], its effects at low doses on gut mucus and bacterial population might improve barrier function. In addition, in IHC analysis, we found that TRPV1 expression in colon sections was significantly decreased in HFD and H+MUC, while capsaicin improved it. TRPV1 expression is reported to decrease in white and brown adipose tissue during obesity [34]. But there are no reports on how its expression profile in gut changes in obesity. Exogenous mucus feeding did not show any improvement on HFD-associated decrease in TRPV1 levels. This was probably because mucus itself or its after-effects could not trigger TRPV1 regulatory pathways in cells. In a separate experiment replicating our previous findings [22] in mice, we saw mucus depletion upon TRPV1 systemic ablation, with downregulation in some of the related genes (Fig.S1, supplementary data). It led us to speculate that TRPV1 modulation by capsaicin must be an essential initiating factor that cascades into mucosal protection and gut barrier reinforcement.

Histological examination of colon sections was performed to see the changes in mucus production. AB stains glycoproteins (specifically glycan parts) and intestinal samples contain mucins as predominant glycoproteins. Besides, HID is used for staining sulfomucins. We found reduced AB intensity in HFD samples, while H+MUC and H+CAP showed an increase in the stain uptake. While changes were not drastic, they were statistically significant. Also, in Carnoy’s solution-fixed samples, we found that HFD reduced inner mucus layer thickness, while mucin and capsaicin improved it. This further reflected the change in mucus production patterns. We expected sulfomucins to be decreased in HFD, similar to total mucins but found an opposite way. Both interventions were able to reverse this effect too. While there is no literature available about how sulfomucins specifically behave in HFD conditions, some studies show that sulfate-reducing bacteria (SRB) are increased in obesity [35], which is related to increased sulfomucins itself. We also found similar patterns in the SRB genus Bilophila. However, no other SRB were changed significantly. An increase in SRB is associated with the onset of IBD and indicated in obesity [36,37]. This finding needs to be explored further.

Nevertheless, exogenous mucins caused effects similar to capsaicin. We could speculate that mucin feeding induced changes in gut microbiota like capsaicin, which further modulated the gut barrier function and mucus production itself. We checked the levels of LPS – an indicator of endotoxemia, and IL-6 – a major reporter of inflammation, in serum samples. Both LPS and IL-6 were increased in HFD and were improved in mucin as well as capsaicin intervention. This further endorsed our hypothesis that mucus modulation was crucially involved in the beneficial effects of capsaicin in enhancing the barrier function and preventing HFD-induced endotoxemia and inflammation. Gene expression analysis in colon tissue showed a downregulation pattern in most genes in the HFD group; however, the expression levels remained mostly unchanged by interventions (Fig.S2, supplementary data). Notably, genes like Tff3, Dll1 (goblet cell maturation), Vamp8 (vesicular transport), Mep1b (mucosal clearance regulation), Reg3g, Def1b (anti-microbial action) showed upregulation by capsaicin treatment.

Besides an SRB genus, in 16S metagenome analysis, we found many bacterial ranks with significant changes. Bray-Curtis ANOSIM showed a closer similarity of H+MUC and H+CAP samples to the Control group than HFD. P-value was 0.001. However, a mid-range R-value (0.674) for overall comparisons indicated relatively high variations among replicates within groups. In the PCA plot, a similar pattern was observed. Upon examining the individual bacterial ranks of interest, the Bacteroidetes-Firmicutes ratio was significantly lower in HFD than in Control. A decrease in the Bacteroidetes-Firmicutes ratio has often been associated with obesity [38], and both mucin and capsaicin reversed it. Lactic acid bacteria are well-established probiotics; they have been associated with barrier function improvement and are known to colonize intestinal mucus [39,40]. Lactobacillus (and some of its reclassified genera – Ligilactobacillus, Limosilactobacillus [41]) and Lactococcus were decreased in the HFD group expectedly. Both interventions prominently improved the abundance of these genera. Lactiplantibacillus (a reclassified *L. plantarum* strain [41]) was increased significantly by both interventions. Clostridium clusters XIVa and XVIII – known butyrate producers and mucus colonizing bacteria [42], were enriched in capsaicin. Schaedlerella – another reclassified arabinose-feeding Clostridium genus [43], and Anaerotignum (reclassified *C. lactatifermentans* [44]) were improved in both mucin and capsaicin administration. Anaerofustis, Fecalimonas, Mediterraneibacter (a reclassified Ruminococcus) and Raoultibacter (recently identified) [45,46] were negatively affected in HFD, which was reversed in both interventions. Pseudoflavonifractor, Intestinimonas are reported to increase in obesity [47,48]. We found similar patterns with significant reversal by both interventions. The genus Sporobacter showed patterns opposite to available literature. Bilophila, an LPS-producing SRB genus, is associated with inflammation and aggravating effects of HFD [49]. We found it increased in the HFD group, though only capsaicin could significantly reduce it. Dysosmobacter, Colidextribacter and Neglecta are some recently identified bacterial genera, with no literature available regarding obesity or other health conditions. We found that HFD altered these significantly and both mucin and capsaicin could revert these effects. More information is required to establish clearly how these genera could associate with obesity. Nonetheless, enough evidence from metagenomic analysis implied that capsaicin-induced changes in microbial diversity were linked to the modulation of gut mucus. While our findings indicate that exogenous mucin could improve gut microbiota like capsaicin, literature also shows that various microbes can affect mucus production itself, both positively and negatively [50]. Therefore, it seems apt to say that mucus and microbes affect each other in a cyclic fashion.

TRPV1 expression is well-established in sensory neurons of DRGs and colonic vagal innervations and non-neuronal cells. Still, there are no reports on TRPV1 expression in mucin secreting goblet cells in the gut. This led us to hypothesize that capsaicin-induced changes in mucus regulation might have neuroendocrine mechanisms instead of a local, direct effect on goblet cells. In addition, while TRPV1 is mentioned as a probable mediator of capsaicin-induced changes in gut microbes, the possibility of direct interactions between capsaicin and bacteria has never been addressed. Some *in vitro* studies report the antibacterial effects of capsaicin [51], but these effects are not considerable at doses corresponding to our study. We tried to understand how gastrointestinal TRPV1 could fit in capsaicin-mucus-microbes interaction and if there was a direct interaction between capsaicin and gut microbes. Using *in vitro* fermentation in fecal samples with capsaicin concentration matching that in the lower gut, we found an increase in some beneficial genera like *Bifidobacterium, Lactobacillus, Akkermansia Eubacteria* and *Fecalibacterium*, however, no significant changes were found in tested Gram negative bacteria. It suggests that capsaicin can influence bacterial diversity *per se*, besides through mucosal modulation or TRPV1-dependent neurotransmitters. Since we noticed enhanced levels of butyrate and increase in the abundance of *Bifidobacterium* and *Lactobacillus* upon capsaicin supplementation, we evaluated whether these agents would stimulate the action of TRPV1 in the transfected HEK cells. Butyrate and live bacteria from all tested strains (Lab3, Lab39, Bif) were able to induce Ca^2+^-influx in transfected cells. Though still in infancy, these findings indicate that certain bacteria and their metabolites can directly modulate TRPV1 activity. Some of the very recent literature also strongly augments our observations in this study [52,53].. This suggests that capsaicin treatment might impede obesity either by direct agonism of TRPV1 or through modulating bacteria or promoting specific bacteria, which could acts as agonists besides microbial metabolites. This study might generate curiosity for TRP researchers in understand the role of selected gut bacteria as agonists for TRP receptors.

Another question is how capsaicin-TRPV1 or capsaicin-bacteria-TRPV1 interactions play into mucus regulation in the gut. Mucus production is regulated in the gut by various signalling pathways. Serotonin, known as a hormone and a neuropeptide, is secreted by EC cells in colon and is hinted to be involved in mucus regulation. However, the information is limited [54]. A recent study has explored the involvement of serotonin in neuro-microbial communication [55]. On the other hand, TRPV1 activation leads to the release of neurotransmitters like CGRP, Substance P, NO etc., which enhance serotonin release and modulate mucus secretion [56,57]. We decided to check Serotonin levels in the colon (primary site of production) and serum (for circulatory levels). HFD expectedly decreased 5-HT in both colon and serum, which both capsaicin and mucin reversed. We also checked serum CGRP levels and found a similar pattern. In both interventions, bacterial diversity seems to be the common factor, causing changes in serotonin levels and mucus production. However, it is difficult to say which element is the primary causative here. Most of the involved factors are modulated by each other, resulting in a cyclic cascade of events rather than a straight pathway.

The evidence in the present study suggests that capsaicin’s anti-obesity effects, besides occurring through augmentation of metabolism and energy expenditure, are in major part via gut barrier and mucosal modulation. Mucus-microbe interactions are crucial to these effects, and TRPV1 must be involved via direct and indirect impacts of capsaicin, provoking neuro-enteric communication. While our findings reveal somewhat limited information on how all these components are interconnected, this study provides new insights into the involvement of TRPV1 and argues for splitting focus towards further exploring mechanisms involved in the enteric effects of capsaicin. These insights will also help study capsaicin or other dietary TRPV1 agonists as a preventive strategy against diseases affecting gut mucus and permeability, such as ulcerative colitis, IBD and Crohn’s disease.

## Author contribution (CRediT author statement)

**Vijay Kumar:** Conceptualization, methodology, investigation, analysis, original draft. **Vibhu Kumar, Neha Mahajan, Jasleen Kaur, Kirti Devi:** Investigation, analysis. **Ravinder Naik Dharavath, Ravindra Pal Singh:** Investigation, analysis, review and editing. **Kanthi Kiran Kondepudi:** Conceptualization, review and editing, resources, supervision. **Mahendra Bishnoi:** Funding acquisition, conceptualization, validation, review and editing, supervision.

## Acknowledgement

Authors would like to thank Department of Biotechnology, Government of India for research grant given to National Agri-Food Biotechnology Institute (NABI) and Dr. Mahendra Bishnoi. Authors thank University Grant Commission (UGC) Government of India for research fellowship given to Mr. Vijay Kumar. Authors are grateful to Dr. Arnab Mukhopadhyay, National Institute of Immunology (NII) for providing facilities and his team for helping to carry out nCounter experiment and analysis.

---

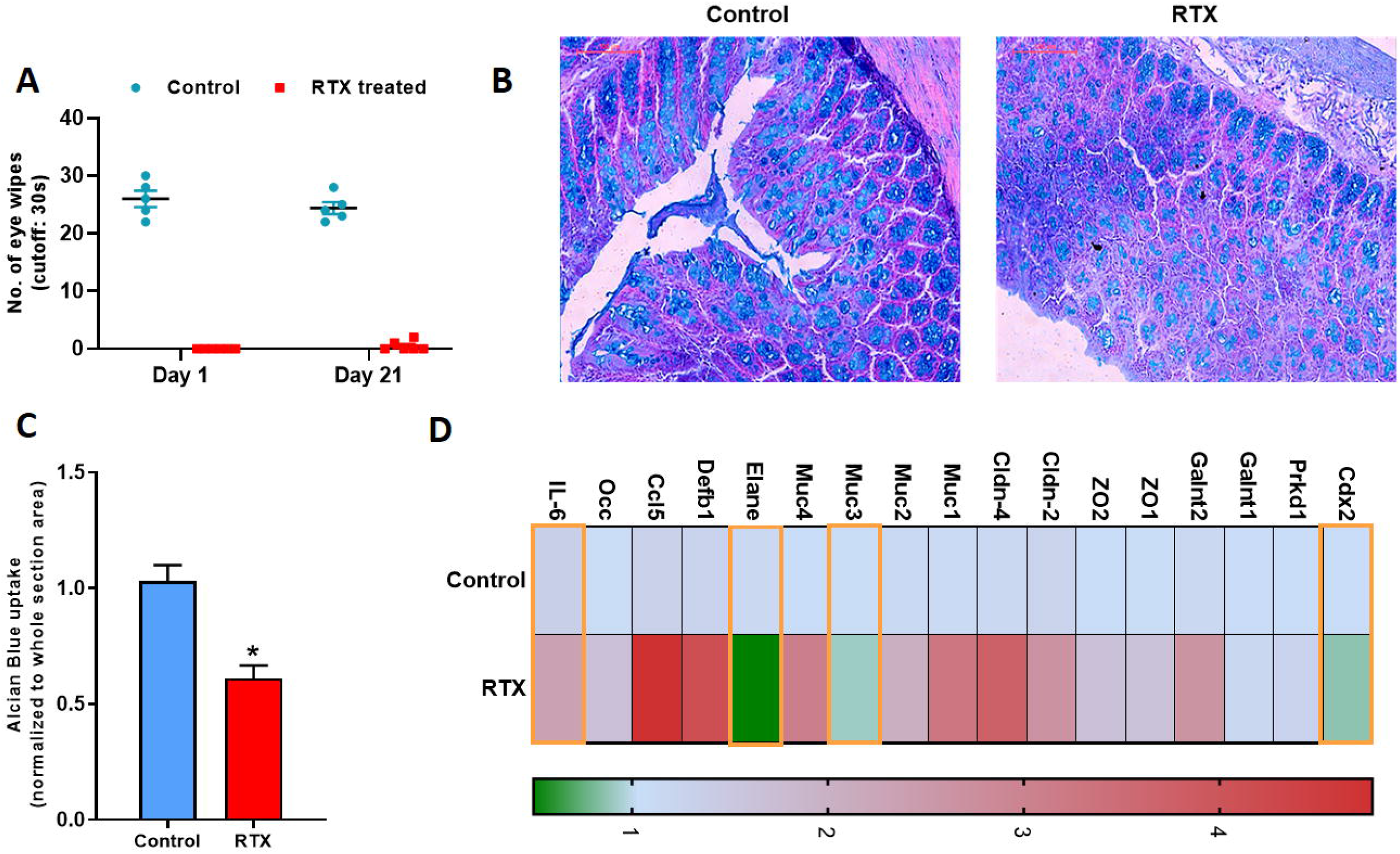

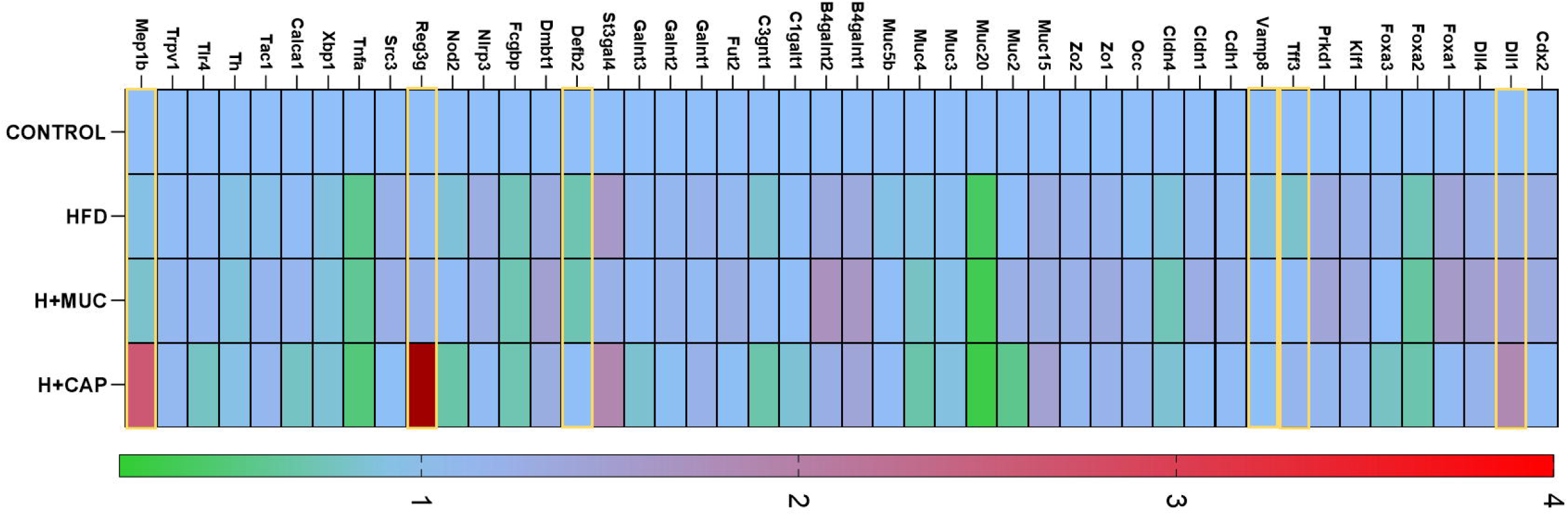

