## Supplementary Data for "Mucin-mimetic action of capsaicin improves high fat diet-induced gut barrier dysfunction in mice colon"

1. **Figure legends**

**Fig.S1. Resiniferatoxin (RTX)-induced TRPV1 ablation.** A) Eye wipe test (capsaicin 0.02%w/v), B) Hematoxylin-Eosin-Alcian blue staining in colon section, C) Alcian blue uptake intensity, D) Heatmap overview of gene expression in colon. Swiss albino mice (n=4 each) were divided into Control (no treatment) and RTX (resiniferatoxin 300µg/kg s.c. injection once). Food and water were given *ad libitum.* Loss of physiological response (eye wipes) was confirmed 24h after treatment and at end of study (after 3 weeks). Animals were sacrificed and decrease in mucus production (histological analysis) was assessed in colon tissues. RNA was isolated from colon samples and gene expression was studied using qPCR. Data is presented as mean±SEM or mean only (heatmap).

**Fig. S2. nCounter data (Heatmap overview of gene expression in colon)**. Mice were divided into 4 groups – Control (normal pellet diet-fed), HFD (high fat diet-fed), H+MUC (HFD + porcine stomach mucin (1g/kg/day p.o.), H+CAP (HFD + capsaicin 2mg/kg/day p.o.). Food and water were given ad libitum. Treatments were given for 12 weeks. After sacrifice, colon tissues were harvested, RNA was isolated and gene expression analysis was performed on commercially customized Nanostring nCounter CodeSet gene expression panel (NanoString Technologies, USA). Data in heatmap is presented as mean values only.

2. **Supplementary Methods:**
a. **Resiniferatxin-induced TRPV1 ablation**

Swiss albino mice (n=4 each) were divided into Control (no treatment) and RTX (resiniferatoxin 300µg/kg s.c. injection once). Food and water were given *ad libitum.* Loss of physiological response (eye wipes test using capsaicin 0.02%w/v) was confirmed 24h after treatment and at end of study after 3 weeks. Animals were sacrificed and decrease in mucus production (histological analysis: see methods section 2.5) was assessed in colon tissues. RNA was isolated from colon samples and gene expression was studied using qPCR.

b. **Gene expression analysis: qPCR**

RT-qPCR was employed for gene expression analysis in colon samples. Total RNA was isolated from tissues using Trizol-Chloroform-Isopropyl alcohol method and quantified on Nanodrop (Thermo Fisher Scientific, Massachusetts, USA). RNA integrity was determined by agarose (1.2%) gel electrophoresis and DNase (Thermo Fisher Sci.) treatment was given to eliminate any genomic DNA contamination. cDNA synthesis was done using RevertAid First Strand cDNA Synthesis Kit (Thermo Fisher Sci.) and relative change in gene expression was determined by qPCR using SsoAdvanced Universal SYBR Green Supermix (Bio-Rad, USA). qPCR was performed on CFX96 Touch Real-Time PCR Detection System (Bio-Rad) under following conditions: initial denaturation – 95°C, 2 min, [denaturation – 95°C, 5 sec; annealing/extension –60°C, 30 sec] x 40 cycles, final extension – 60°C, 5 min and melt curve analysis between 60°C-95°C with 0.5°C/5 sec increment. Data was analyzed using ΔΔCt method [1], β-actin was used for normalization. The list of primers used in the experiment is given in Table 2 (supplementary data).

c. **Gene expression analysis: nCounter**

A commercially available Nanostring nCounter CodeSet gene expression panel (NanoString Technologies, USA), comprising of 46 probed genes associated with goblet cell differentiation, tight junction proteins, mucin production, glycosylation enzymes, immune response, peptidergic neuronal markers with 4 housekeeping genes was employed to generate gene expression data. Briefly, total RNA was extracted from colon samples using TRIzol RNA extraction reagent-based method. 50ng sample was hybridized with unique probes to the target RNAs in nCounter CodeSet gene expression panel using NanoString nCounter prep station and placed into the cartridge (NanoString Technologies, USA). NanoString nCounter digital analyzer (NanoString Technologies, USA) was employed for detection and counting of hybridized probes. The data was analyzed using nSolver 4.0 software (NanoString Technologies, USA). All the procedures were performed following the NanoString guidelines.

**Table 1.List of primers used for qPCR in bacterial abundance estimation.**

| **No.** | **Primer ID** | **Forward primer 5'-3'** | **Reverse primer 5'-3'** |
| --- | --- | --- | --- |
| 1 | **Total Bacteria_d** | GCAGGCCTAACACATGCAAGTC | CTGCTGCCTCCCGTAGGAGT |
| 2 | **Bacteroidetes_p** | ACGCTAGCTACAGGCTTAACA | ACGCTACTTGGCTGGTTCA |
| 3 | **Firmicutes_p** | GCGTGAGTGAAGAAGT | CTACGCTCCCTTTACAC |
| 4 | **Akkermensia_g** | AACGAACGCTGGCGGCGTGGATAAGACAT | CATCCCAGTTACCAGTCTCACCTTAGGACCCT |
| 5 | **Bacteroides_g** | GAGAGGAAGGTCCCCCAC | CGCKACTTGGCTGGTTCAG |
| 6 | **Bifidobacterium_g** | GATTCTGGCTCAGGATGA | CTGATAGGACGCGACCCC |
| 7 | **Cronobacter_g** | TGTCTGGGAAACTGCCTG | TCTCAGACCAGCTAGGGA |
| 8 | **Eubacteria_g** | GCTGTGAAGCCGAGCAAA | GGTTAGGTCACTGGCTTC |
| 9 | **Fecalibacterium_g** | GAGGAAGATAATGACGGTAC | ACCTCTGCACTACTCAAGA |
| 10 | **Fusobacterium_g** | CCCTTCAGTGCCGCAGT | GTCGCAGGATGTCAAGAC |
| 11 | **Lactobacillus_g** | CACCGCTACACATGGAG | AGCAGTAGGGAATCTTCCA |
| 12 | **Prevotella_g** | GGTGTCGGCTTAAGTGCCAT | CGGAYGTAAGGGCCGTGC |
| 14 | **Vibrio_g** | CTGCGTCCCGTAGGAGTTTG | ACAGGGGGATAGCAGTTGGA |

_d = domain, _p = phylum, _g = genus.

**Table 2.List of primers used for qPCR in RTX-induced TRPV1 ablation experiment.**

| **No.** | **Primer ID** | **Forward primer 5'-3'** | **Reverse primer 5'-3'** |
| --- | --- | --- | --- |
| 1 | **B-act** | TGTTACCAACTGGGACGACA | GGGGTGTTGAAGGTCTCAAA |
| 2 | **Cdx2** | GCAGAGCCAAGGAGAGGAAA | CTTGCAAGGAGGTCACAGGA |
| 3 | **Prkd1** | CACCCAACCCGTGGAAGG | ACCTGGAGTCATCGCTTTCG |
| 4 | **Galnt1** | GGGAAACCAGTCGTCATTCC | GTGCTCCATGCCTCATTGTG |
| 5 | **Galnt2** | CTGGGCATCGCCTACTACAT | GCCTCCTGGTTAAAGTCTGG |
| 6 | **ZO1** | TAACTTGGGGAGGGAGGGTC | GGTAAGGCATTCCTGCTGGT |
| 7 | **ZO2** | CCCAGAATGCGAGGATCGAA | CTGGAAGGAGCTTTCTGGGG |
| 8 | **Cldn-2** | TCACACTTGAGTCATCGCCC | TCCCACCTCAAGCACAATCC |
| 9 | **Cldn-4** | GACCTAGAAGCAGCCCAGTG | CTCAGAGGGGCCAACTCAAG |
| 10 | **Muc1** | GCCGAAAGAGCTATGGGCA | CTGCCATTACCTGCCGAAAC |
| 11 | **Muc2** | GGCCTCACCACCAAGCGTCC | TGGGCTGGCAGGTGGGTTCT |
| 12 | **Muc3** | AGTGCTGTTGGTGATCCTCG | AGAGTCCAGGGGCATGTAGT |
| 13 | **Muc4** | TTGCACCTGTCCCCCCTGCCT | GTTCGCCACCGAGGCGTTGA |
| 14 | **Elane** | ACCATCACCCAGTGTGCTAC | GCGAAGGCATCTGGGTACAA |
| 15 | **Defb1** | TCCCAGATGGAGCCAGGT | AGCTGGAGCGGAGACAGA |
| 16 | **Ccl5** | ATCATCCTCACTGCAGCCG | TTCTCTGGGTTGGCACACAC |
| 17 | **Occ** | GGCAAGCGATCATACCCAGA | TCATAGTGGTCAGGGTCCGT |
| 18 | **IL-6** | GATGGATGCTACCAAACTGGA | GAGCATTGGAAATTGGGGTA |

**Abbreviations:** B-act = β-actin, Cdx2 = Caudal type homeobox 2, Prkd1 = Protein kinase D1, Galnt1/2 = GalNAc transferase 1/2, ZO1/2 = Zona occludens 1/2, Cldn-2/4 = Claudin 2/4, Muc1/2/3/4 = Mucin 1/2/3/4, Elane = Elastase, Neutrophil Expressed, Defb1 = Defensin-β1, Ccl5 = Chemokine ligand 5, Occ = Occludin, IL-6 = Interleukin 6.
